# Quantifying the Confidence in fMRI-based Language Lateralisation through Laterality Index Deconstruction

**DOI:** 10.1101/530626

**Authors:** Martin Wegrzyn, Markus Mertens, Christian G. Bien, Friedrich G. Woermann, Kirsten Labudda

## Abstract

In epilepsy patients, language lateralisation is an important part of the presurgical diagnostic process. Using task-based fMRI, language lateralisation can be determined by visual inspection of activity patterns or by quantifying the difference in left- and right-hemisphere activity using variations of a basic formula [(L-R)/(L+R)]. However, the values of this laterality index (LI) depend on the choice of activity thresholds and regions of interest. The diagnostic utility of the LI also depends on how its continuous values are translated into categorical decisions about a patient’s language lateralisation. Here, we analysed fMRI data from 712 epilepsy patients who performed a verbal fluency task. Each fMRI data set was evaluated by a trained human rater as depicting left-sided, right-sided, or bilateral lateralisation or as being inconclusive. We used data-driven methods to define the activity thresholds and regions of interest used for LI computation and to define a classification scheme that allowed us to translate the LI values into categorical decisions. By deconstructing the LI into measures of laterality (L-R) and strength (L+R), we also modelled the relationship between activation strength and conclusiveness of a data set. In a held-out data set, predictions reached 91% correct when using only conclusive data and 82% when inconclusive data were included. Although only trained on human evaluations of fMRIs, the approach generalised to the prediction of language Wada test results, allowing for significant above-chance accuracies. Compared against different existing methods of LI-computation, our approach improved the identification of inconclusive cases and increased the accuracy with which decisions for the remaining data could be made. We discuss how this approach can support clinicians in assessing fMRI data on a single-case level, deciding whether lateralisation can be determined with sufficient certainty or whether additional information is needed.

## Introduction

In patients with refractory focal epilepsies, brain surgery is recommended as an effective treatment option (Kwan et al., 2009). To plan such an intervention, presurgical diagnostics aim to identify both the epileptogenic cortex and eloquent parts of the cortex that need to be spared in order to avoid cognitive deficits (Rosenow, 2001; Labudda et al., 2010). Language is of vital importance for everyday functioning and is usually strongly lateralised to one hemisphere of the brain (Knecht, 2000). In such a case, unilateral resection of eloquent cortex cannot be well-compensated by the contralateral homologue, making language lateralisation one key objective of presurgical diagnostics (Binder et al., 1996; Sabsevitz et al., 2003). While most people in the general population show left-lateralised language functions (Springer et al., 1999), the ratio of atypically lateralised cases is higher in epilepsy patients. It is estimated that around 20% of the patients show bilateral or right-sided lateralisation of language functions (Springer et al., 1999). For brain surgery to be safe and beneficial, the certainty regarding the patient’s language lateralisation needs to be maximised.

To determine language lateralisation, both invasive and non-invasive methods can be used (Binder et al., 1996; Rutten et al., 2002; Woermann et al., 2003). Among the non-invasive methods, functional MRI is recommended as a reliable diagnostic tool for the lateralisation of language functions (Binder, 2011; Szaflarski et al., 2017). A frequently used paradigm to estimate language lateralisation with fMRI is verbal fluency, a task in which the patient silently generates as many words as possible belonging to a certain category (Henry and Crawford, 2004). The words to be generated need to belong to either a semantic category (for example, fruits or animals) or a phonological category (for example, words beginning with S). A verbal fluency task mainly measures language production, activating regions in the inferior frontal gyrus (IFG), including Broca’s area (Gaillard et al., 2000; Arora et al., 2009). Depending on the broadness of the comparison condition, other areas such as the supplementary motor area (SMA), visual word form area (VWFA) in the fusiform gyrus, and Wernicke’s area will also show activity, giving rise to a distributed but lateralised language network (Lurito et al., 2000; Price, 2010).

One way to determine the language lateralisation of a patient based on fMRI results is to compute a single laterality index (LI) from the emerging voxel-wise activity patterns. This index indicates the difference in language-related activity in the left versus the right hemisphere (Chlebus et al., 2006; Jansen et al., 2006). Such an index is usually computed by counting the voxels in each hemisphere’s language-related areas falling above a predefined activity threshold. This approach requires the evaluator to decide which activity threshold and regions of interest (ROIs) to evaluate (Gaillard et al., 2002; Adcock et al., 2003). Once the number of above-threshold voxels in a certain brain area has been computed, the difference between the left and right side is expressed in a single value. The most common formula used is (L-R)/(L+R), which gives the difference between the above-threshold voxels in a region of interest in the left and right hemispheres divided by the sum of the above-threshold voxels in both hemispheres (Seghier, 2008).

Finally, the resulting score must be translated into a categorical decision by using a cut-off so that, for example, cases with LI values above +0.2 will be categorised as left-lateralised, cases below −0.2 as right-lateralised, and cases in between as bilateral (Wilke et al., 2011). Defining these cutoffs is difficult, partly because the sample sizes of validation studies tend to be small (Dym et al., 2011). For example, if a sample includes only one atypical case with an LI value close to −1, a wide range of cutoffs can produce perfect accuracies.

Nevertheless, many approaches based on variants of laterality indices have demonstrated high concordance with invasive measures (Jones et al., 2011; Janecek et al., 2013) or language-related clinical variables such as handedness or age at onset (Berl et al., 2014). Still, the common LI has been criticised because it is threshold-dependent and ignores the high inter-individual variability of fMRI activity strength (Suarez et al., 2009; Strandberg et al., 2010).

For instance, in a patient with very poor activity, nine above-threshold voxels in the right hemisphere and one above-threshold voxel in the left hemisphere might remain after applying a moderate threshold. The resulting LI would be (1-9)/(1+9) = −0.8, indicating atypical, right-sided language dominance. In contrast, the underlying fMRI activity pattern would most likely be considered inconclusive by a human expert, as the overall signal would be insufficient to draw conclusions about the functional organisation of language. Hence, a fixed threshold LI will assign the most extreme scores to the patients with the fewest above-threshold voxels (i.e., with the poorest data quality). To address this problem, more sophisticated methods use variations of adaptive thresholds (Wilke and Lidzba, 2007). There, weaker activations are thresholded at lower levels, ensuring that the LI computation is always based on an adequate amount of data in the individual case. However, if an adaptive threshold allows more random noise to pass the lower threshold, the difference between left and right hemisphere will diminish, and poor-quality data will be reflected in LI values closer to zero, indicating bilaterality. Because of these issues, methods for LI computation critically rely on data pre-selection (Wilke and Lidzba, 2007) usually based on a subjective criterion. In summary, the reliability of analysing a language-fMRI arguably depends upon (i) deciding how to compute the LI, (ii) from which regions of the brain to extract data, (iii) how to translate the continuous LI values into categories of lateralisation, and (iv) how to decide which data sets do not allow for making a decision with sufficient confidence.

In the present study, we aimed to evaluate how the fMRI activity patterns from a verbal fluency task can be best used for assessing the type of language lateralisation of a patient with epilepsy. We used the common LI [(L-R)/(L+R)] applied to different activity thresholds and ROIs. We trained a classifier to determine the cutoffs that best allow for grouping the continuous LI values into categories of lateralisation. These categories were based on a trained specialist’s free inspection of the fMRI data (Woermann et al., 2003). Each data set was categorised as indicating left, bilateral, or right language lateralisation or as being inconclusive (i.e., refraining from making a decision). To improve the identification of inconclusive data, we deconstructed the LI formula into a measure of lateralisation (L-R, its numerator) and activity strength (L+R, its denominator). This was aimed at addressing the ambiguity of the single-value LI discussed above, especially in the case of low-quality data.

To compare the performance of our approach against a benchmark, we used already established methods of LI computation (Wilke and Schmithorst, 2006; Wilke and Lidzba, 2007) as a frame of reference. Finally, we validated our approach using language Wada test results as the gold standard for language lateralisation.

## Methods

### 2.1 Participants

The study included fMRI data from 712 patients with epilepsy who were undergoing a presurgical evaluation. All patients performed an fMRI verbal-fluency task at the Epilepsy Centre Bethel between September 2011 and March 2018. The start of the inclusion period was determined by the installation of a 3T scanner at the study site. All data were acquired as part of the centre’s presurgical evaluation programme, and data were analysed retrospectively. The study was approved by the ethics board of Bielefeld University (2017-184). The full data set for the present study comprised 783 fMRI sessions, as some patients were scanned on multiple occasions. Of the included patients, 46% were female, the median age was 28 years (range: 4–74), and 81% were right-handed.

### 2.2 MRI data acquisition

Data were collected on a 3T Siemens Verio MR scanner. High-resolution T1-weighted structural data were collected for each patient using a 32-channel head coil with 192 sagittal slices, slice thickness of 0.8 mm, and 0.75 x 0.75 mm in-plane resolution. For fMRI data, a 12-channel head coil was used, and data were collected using the following parameters: 21 axial slices per volume, 3×3 mm in-plane resolution for each slice, and a thickness of 5 mm. A repetition time (TR) of 3 seconds was used with an effective acquisition time of 1.8 seconds per volume; there was a 1.2-second pause between TRs to allow for the audible presentation of verbal instructions. Over a period of 10 minutes, 200 volumes (excluding two dummy scans) were collected.

### 2.3 fMRI task

The verbal fluency task consisted of covert word production of either semantic or phonemic categories such as “animals” or words beginning with the letter “S.” A block of verbal fluency lasted 30 seconds, and the blocks were alternated with a 30-second resting condition when the patients were asked to stop generating words and relax. There were 10 blocks for each condition, each triggered by a verbal instruction given via the MRI intercom, resulting in a task length of 10 minutes. If necessary, patients were trained to perform the task beforehand by a neuropsychologist, with some patients receiving specifically tailored lists of categories in accordance to their abilities and interests.

### 2.4 Visual evaluation

To provide a reference for the evaluation of the LIs, the assessment by a trained specialist (FGW) for each case was derived from clinical records. This assessment was based on an unrestrained visual inspection of the whole-brain activity patterns, which included varying the threshold at which the activity distribution was deemed meaningful. Visual inspection included an assessment of data quality; stimulus-associated movements (often present along tissue borders) or the absence of a default mode network activation pattern during rest were used to identify low-quality data. Also, the specificity of the activity patterns was considered so that more weight was given to a pattern that encompassed the IFG, SMA, and VWFA as opposed to a pattern of activity that was restricted to the precentral gyrus, which might be due merely to the patient co-moving their lips. Each fMRI was coded as either left-hemispheric, right-hemispheric, bilateral, or inconclusive language lateralisation. Accordingly, the sample consisted of 527 left-lateralised (67%), 75 bilateral (10%), 47 right-lateralised (6%), and 134 inconclusive (17%) cases. Excluding the inconclusive cases, the distribution of lateralisation was 81% left, 12% bilateral, and 7% right.

### 2.5 Data preprocessing

For statistical analyses, data were preprocessed using SPM12 with the following steps: First, the fMRI time series was movement corrected using the realignment function. Then, the images were directly normalised to the echo-planar imaging (EPI) template, up-sampled to 2 mm isotropic voxels, and smoothed with a Gaussian kernel of 6mm full-width at half-maximum (FWHM) to improve the signal-to-noise ratio. This approach was chosen over deriving the normalisation parameters from the co-registered structural scans, as the direct transformation of EPI images proved more robust for many patients, especially when lesions or signal dropouts were present (Calhoun et al., 2017).

### 2.6 Generation of statistical maps

Voxel-wise whole-brain analyses were performed for each patient using SPM12. The block-wise activity was modelled with a canonical hemodynamic response function (HRF). Movement parameters were included as regressors of no interest. A map of t-values was then computed for the comparison of “verbal fluency > rest,” and these maps were used in the subsequent steps to compute laterality indices.

### 2.7 Study design

To arrive at unbiased estimates of accuracy, we split the sample into a training and a testing set, with two thirds of the data used for training and one third held out for testing. As the distribution of classes was uneven (i.e., there were many more left-lateralised patients than atypical patients), we used stratified splitting, so that the base rate for each class was preserved in both splits. Different parameters of t-value thresholds and ROI sizes were used on the training data. Only after training, the best parameter combinations were used to predict the held-out testing data set of 262 fMRI sessions. We used a linear support vector classifier (SVC), which allowed for making probabilistic predictions. This means that the probability of a data set belonging to each class of lateralisation can be expressed as a value between 0 and 1, which allows the degree of certainty to be assessed for each decision (Wu et al., 2004). The splitting and classification of data was implemented using the free software library scikit-learn 0.18 (Pedregosa et al., 2011) in Python 2.7. Code underlying all analyses is available at github.com/mwegrzyn/laterality-index-deconstruction.

### 2.8 Group analysis and definition of regions of interest

The training data were used to compute maps of average activity for each class of language lateralisation as defined by the human evaluator (left, bilateral, right, or inconclusive). Whole-brain one-sample t-tests for each of the four groups were performed using nilearn 0.3 (Abraham et al., 2014) in Python 2.7. Furthermore, the group results for the left- and right-lateralised cases, as defined by the human evaluator, were used to create ROIs. To arrive at a representative map of differences between the dominant and non-dominant hemispheres, the data were transformed as follows. First, the right-lateralised cases were flipped from left to right so that the dominant hemisphere was on the left for all patients. Then, for all of these images, another flipped image was created and subtracted from the original, resulting in images with a left-right difference value for each voxel. Subsequently, the left-right difference images of all patients were used to compute a whole-brain one-sample t-test, resulting in a map where each voxel’s positive t-value on the left side indicated stronger lateralisation to the left. This map of the left hemisphere was then turned into binary masks at different levels of activity. We generated 20 binary maps, starting with all voxels of the map and subsequently dropping 5% of the lowest scoring voxels until only the top 5% of the original map (i.e., the most significant values) remained. To generate ROIs for the right hemisphere, the map was flipped along the x-axis, providing a mirror-image of all regions on their contralateral homologue of the brain.

### 2.9 Computation and deconstruction of the LI

From each patient’s t-map, the values of all voxels inside the left and right language ROIs were extracted. From the lowest to the highest t-values present in the brain map, the percentage of above-threshold voxels was determined –that is, starting at the point when all voxels crossed the respective threshold up to the point where no voxels crossed the threshold. Next, the LI, defined as (L-R)/(L+R), was computed. Each LI can be broken down into its numerator (L-R), which reaches its maximum when the absolute difference in the above-threshold voxel is highest, and the denominator (L+R), which decreases as the threshold increases. To take full advantage of the information contained within the LI, we explored a two-dimensional approach that uses L-R (the numerator) as a measure of laterality and L+R (the denominator) as a measure of activation strength (Fig. 1).

**Figure 1.**
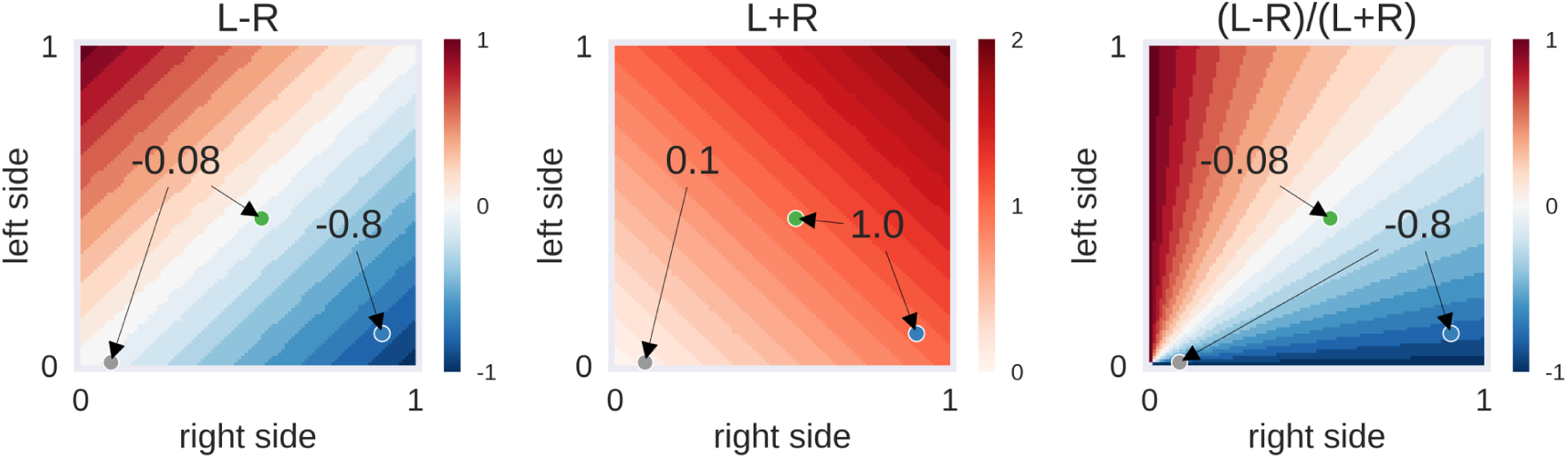
Example of laterality scores depending on the proportion of above-threshold voxels. L-R: the behaviour of the numerator of the laterality index (LI), depending on the proportion of above-threshold voxels in the left and right hemisphere. L+R: the behaviour of the denominator of the LI. (L-R)/(L+R): the LI itself, i.e., the ratio of numerator and denominator. Three hypothetical cases are presented as dots: The grey case has a low voxel count in both hemispheres (L=0.01, R=0.09), the green case has a moderate voxel count in both hemispheres (L=0.46, R=0.54) and the blue case has a high voxel count in the right hemisphere only (L=0.10, R=0.90). While the grey and green case are identical in their left-right difference (L-R), the grey case differs from the others in its low overall activity (L+R). This in turn makes its LI score identical to the blue case. Depending on the approach used, one could decide to label the activity of the grey case as being suggestive of bilaterality (L-R is low) or of strong right-lateralisation (LI is high). The former approach would be taken by adaptive thresholding methods while the latter approach would be employed by fixed-threshold methods. However, another possibility could be to use the information about the strength of overall activity (L+R is low) to classify this case as inconclusive.

### 2.10 LI-Toolbox as benchmark

To compare our approach against a benchmark, we computed laterality indices using the LI-Toolbox (Wilke and Lidzba, 2007) as implemented in SPM12. To be comprehensive, we used all of the following methods for LI computation: (i) a fixed-threshold method at the default value of t=3 where the LI is based on the voxel count; (ii) an adaptive method in which the threshold is set at the mean intensity of the voxels; and (iii) a bootstrap method with its overall LI. For all variations of the LI we used standard settings as recommended in Wilke and Lidzba (2007) and Wilke and Schmithorst (2006). A frontal lobe mask with midline exclusion served as the ROI. Corresponding to the approach outlined in section 2.7, the different toolbox-based LI values of all patients from the training set were used to construct linear SVCs, which were then used to predict the labels of the test data.

### 2.11 Validation with Wada test results

#### 2.11.1 Wada test procedure

Wada testing was performed by internal carotid artery injection of 200 mg amobarbital via a transfemoral catheter separately for each hemisphere. For children, size adapted doses were administered (100mg to 150 mg). Before the intracarotid amobarbital procedure (IAP), cerebral angiography was performed. The IAP language test protocol assessed seven language functions: (1) series repetition (counting); (2) following verbal commands by pointing to an image (four tasks); (3) following body commands (two tasks); (4) visual confrontation naming (four tasks); (5) repetition of sentences or proverbs (two tasks); (6) reading sentences aloud (two tasks); and (7) spontaneous speech. For each of the seven functions, a score of 2 points could be reached.

#### 2.11.2 Categorisation of Wada test results

For bilateral Wada tests (n=44), lateralisation indices were calculated as LI=[(L-R)/(L+R)]*(n/m) (n: score of best hemisphere; m: highest possible score; Kurthen et al. 1994). For unilateral Wada tests (n=21), the hemispheric language capacity was calculated as HLC_h_=h/m (h: score of tested hemisphere, m: highest possible score; Wellmer et al. 2005). The distribution of the LIs and HLCs is illustrated in Fig. 2. The LI scores were categorised as follows: LI>0.5: left-sided dominant; LI<0.5 right-sided dominant; −0.5≥LI≥0.5: bilateral. This follows the categorisation of Kurthen et al. (1994), treating cases showing “incomplete” lateralisation as lateralised instead of bilateral. The HLC_h_ scores were categorised as follows: HLC_h_=0: the tested hemisphere h is dominant; HLC_h_≥0.8: the tested hemisphere is not dominant; 0<HLC_h_<0.2: patients in this range (n=3) were excluded from further analyses because of the possibility of negative bilaterality (Wellmer et al., 2005).

**Figure 2.**
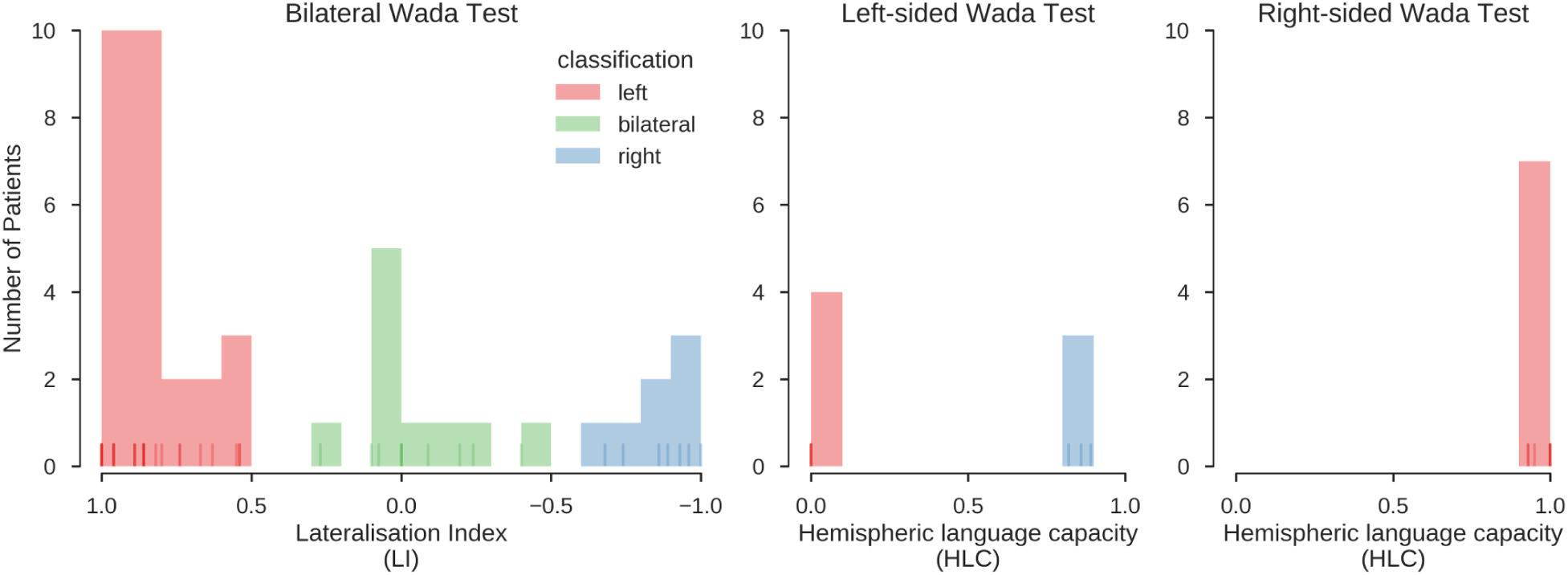
Distribution of Wada test scores. Results for the bilateral and unilateral Wada tests. Histograms indicate the number of cases with a Wada test score falling into the respective bin. Lines at the bottom of the plots indicate individual values (overlapping values indicated by stronger hue). Red, green and blue colours indicate the classification of the scores into left, bilateral and right, using the procedures described in Kurthen et al. (1994) and Wellmer et al. (2005). In the final sample, 39 patients were left-lateralised (63%), 10 were bilateral (16%) and 13 were right-lateralised (21%). This classification was also confirmed using an unsupervised clustering method (K-means clustering, searching for a three-cluster solution), which grouped all data in the same way.

#### 2.11.3 Prediction of Wada test results

The same classifiers that were used to predict the human evaluations were used to predict the result of the Wada test (left, bilateral, right) from the fMRI data. This means that all Wada test results were only used for testing, but not to train the classifier. Also, the fMRI data of all patients with Wada test results were only part of the test set (see section 2.7 study design), and no data from patients with Wada test results were used during training. The predictions of the LI-Toolbox, computed as described in section 2.10, were also validated with the Wada test results using the same procedures.

## Results

### 3.1 Analyses at the group level

The whole-brain activity patterns for the training data are shown in Fig. 3. At the group level, a clear activation pattern emerged including IFG, fusiform gyrus, and SMA. There was also activity in the thalamus on the dominant side and the contralateral hemisphere of the cerebellum. This pattern held true for both left-and right-lateralised cases. The same set of regions was activated for the bilateral cases, although without signs of hemispheric differences, as would be expected. The inconclusive cases showed only a small above-threshold cluster in the SMA and some indication of frontal activity. In addition, all groups showed more activity in the precuneus and orbitofrontal areas during rest and in the superior temporal and angular gyri. This indicates engagement of the default mode network during periods of rest. Of note, supra-threshold activity in Wernicke’s area during verbal fluency was lacking in all groups.

**Figure 3.**
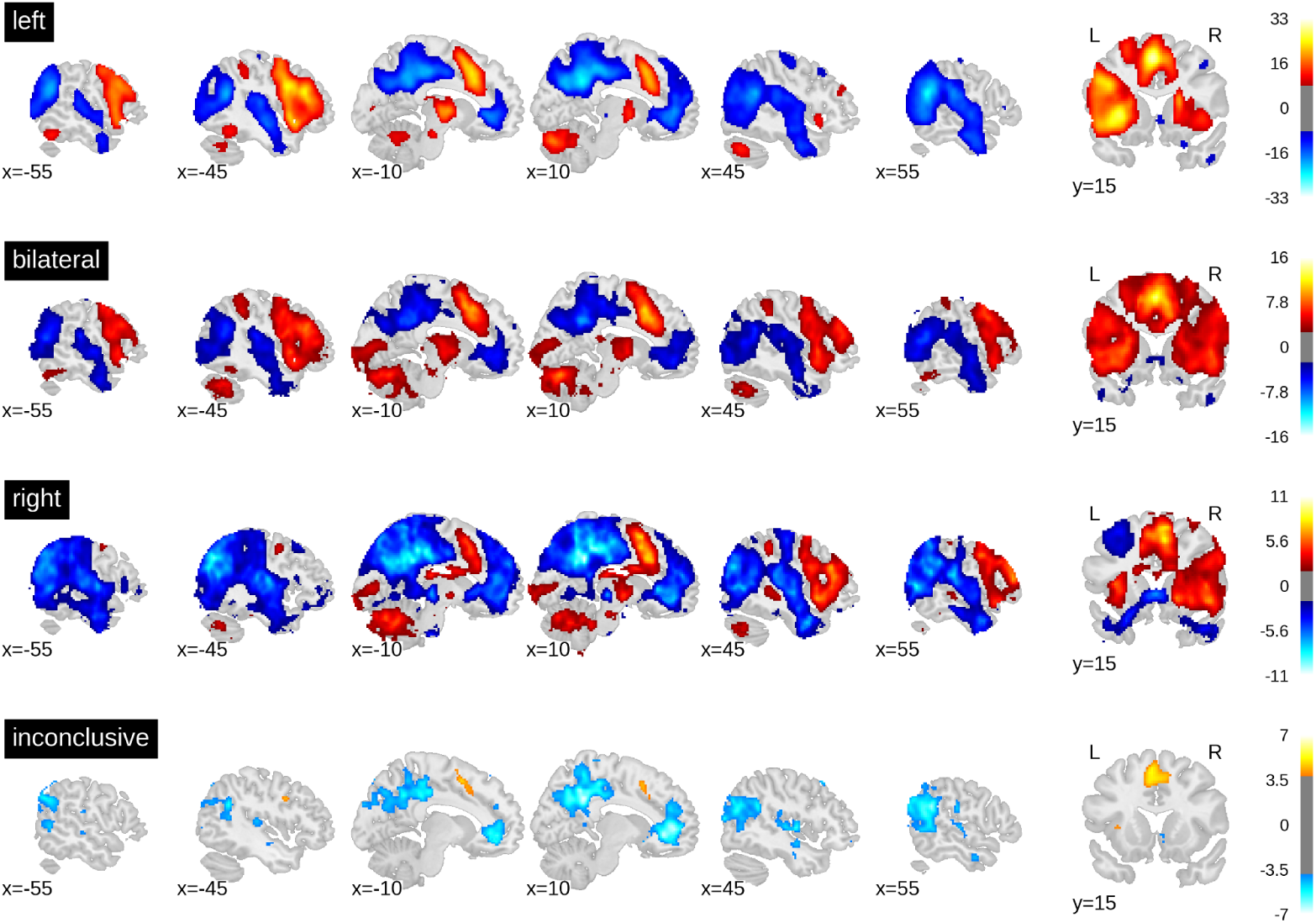
t-value maps from the group level whole-brain analyses. Higher activity for the task is plotted in red and higher activity during rest is plotted in blue. As groups are of different sizes, the threshold for each map has been adjusted for the respective sample size. The required alpha level for a medium effect (d=0.5) and 80% power was computed using GPower 2.1. For the left-sided group (n=351) the critical t is equal to 8.47. For the bilateral group (n=50), t=2.68. For the right-lateralised group (n=31), t=1.93. For the inconclusive group (n=89), t=3.85. Bars on the right-hand side show colour-coding of t-values. Full unthresholded maps are available online: neurovault.org/collections/3887.

When computing a pattern of differences between each voxel in the dominant vs. non-dominant hemisphere and when setting a threshold to include all voxels with t-values greater than zero, the result was an ROI that included all grey matter regions except those associated with the default mode network. When only the top 5% of voxels with highest t-values were included (i.e., the 95th percentile), only frontal areas and part of the fusiform gyrus constituted the ROI (Fig. 4).

**Figure 4.**
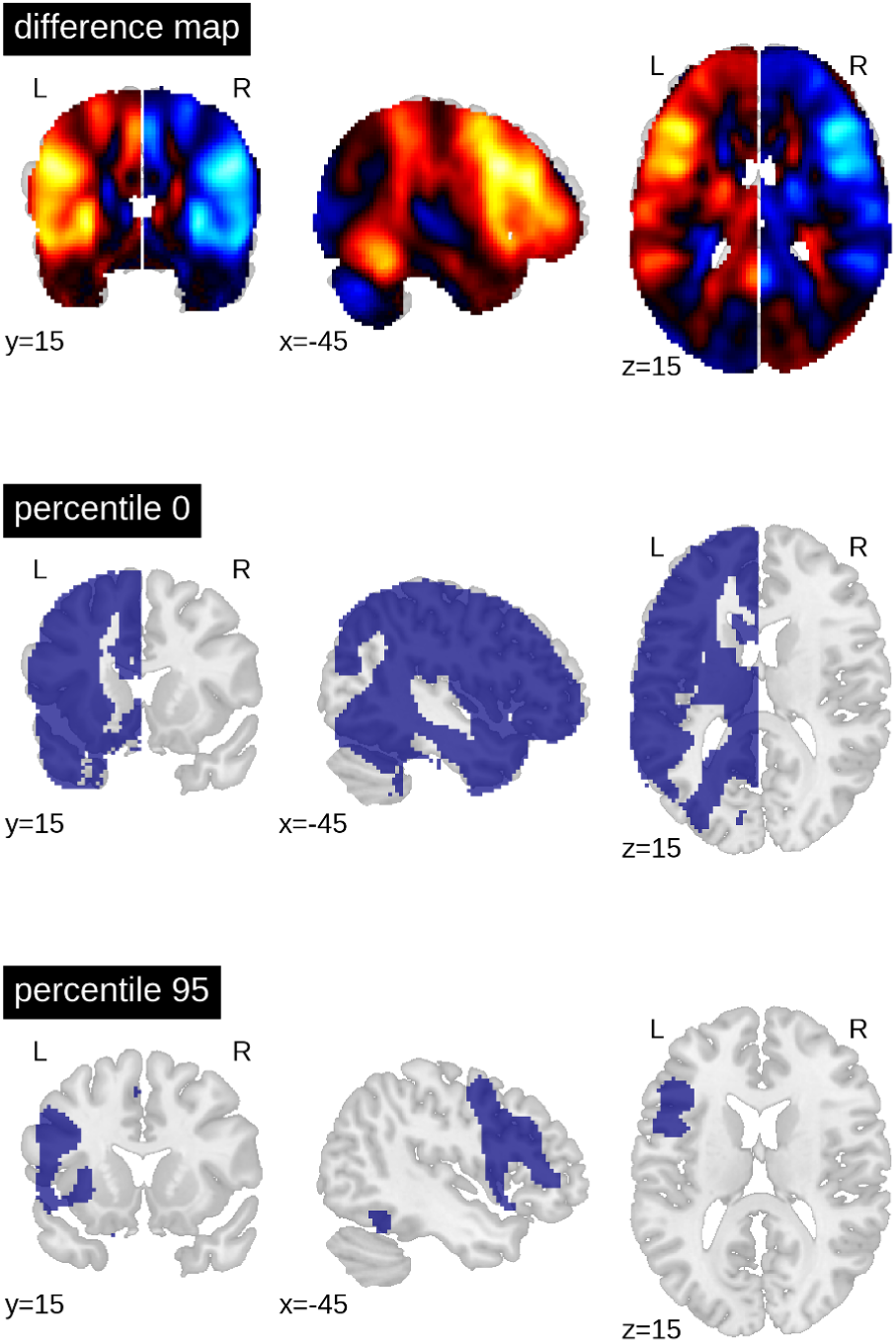
Generation of ROIs. The top brain map shows the results of the one-sample t-test of all voxels in the dominant (left) vs non-dominant (right) hemisphere against zero (for all patients with left- and (mirrored) right-sided lateralisation). The lower maps show example ROIs when including all voxels in the dominant hemisphere larger than zero or when including only the top 5% (95th percentile). The difference map is available online: neurovault.org/images/113673.

### 3.2 Finding optimal parameters

For each combination of activity threshold and ROI size, the training data set was used to compute the accuracy with which the different lateralisation categories could be predicted. To this end, the training data (n=521) were split randomly into two halves with one half used to train the SVC and the other half to compute its accuracy. This nested cross-validation was performed for 100 random splits of data, and the average accuracy was computed for each combination of parameters. This gave rise to a heatmap of accuracies (Fig. 5) across different thresholds of t-values (x-axis) and ROI sizes (y-axis).

**Figure 5.**
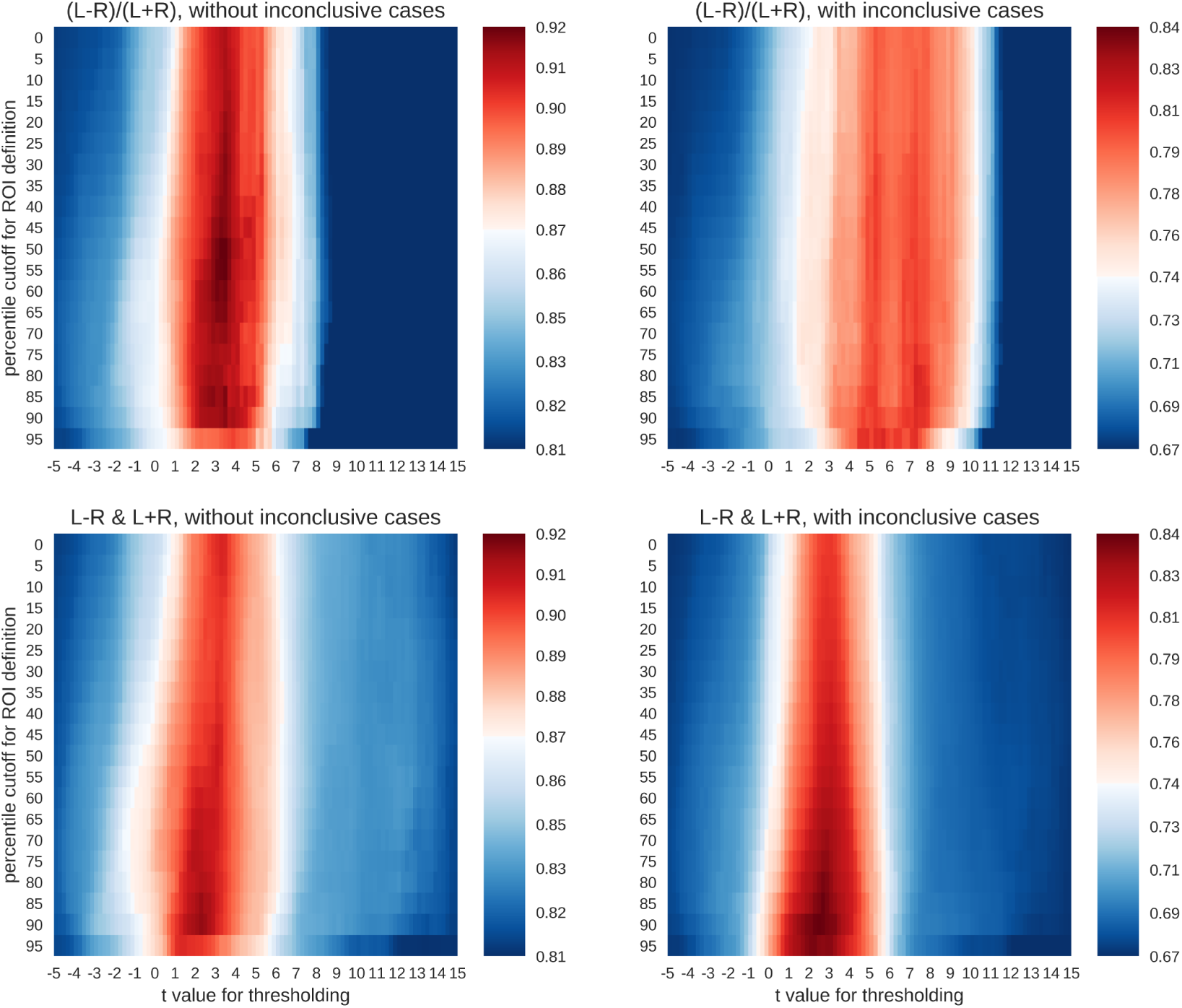
Accuracy maps for different classification rationales. The maps show colour-coded accuracies for different t-thresholds (x-axis) and ROI sizes (y-axis). Predictions were based on random splits of the training data set into two halves. The lowest accuracies are plotted in dark blue and the highest in dark red. The red colour indicates that an accuracy score is significantly better than guessing at p<.001.

To evaluate whether the accuracies were larger than expected by chance, a binomial test was carried out to test the accuracy of each parameter combination against guessing (alpha=.001). The guessing rate (defined as the base rate of the largest group) for the three-class case was 81%, so that above-chance accuracies must reach at least 87% to be considered meaningful. On the other hand, when all four classes were included, the guessing rate dropped to 67%, and the above-chance accuracies must reach at least 74% to be meaningful. As shown in Fig. 5, both the common LI and the two-dimensional approach with laterality and strength allowed for above-chance predictions. This held true when only conclusive cases were included in the analyses or if all data were used. The LI and the two-dimensional approach reached accuracies of 92% and 91% respectively for the three-class case and accuracies of 81% and 84% respectively for the four-class case. However, the two approaches differed in the parameter space that allowed for robust above-chance classification. When only conclusive data were used, both approaches needed t-value thresholds around three to allow for successful classification. However, the common LI worked well for a wide range of ROI sizes, while the two-dimensional approach required more circumscribed ROIs. When inconclusive data were included, the t-value thresholds required by the two approaches differed more prominently. The common LI needed higher thresholds with t-values in the range of three to nine, while the two-dimensional approach did not require a raise in the threshold and still reached its highest accuracies for t-values around two or three. To better understand the differential behaviour of the approaches, we first evaluated the confusion matrices for the top parameter combinations for each approach and then plotted the underlying raw data.

### 3.3 Prediction using the test data

To evaluate how well new data could be predicted and what kinds of mistakes were made in doing so, all above-chance parameters from the computed threshold x ROI maps were used on the held-out testing data set of 262 fMRI sessions. This means that for each held-out patient’s data, the predictions from all above-chance parameter combinations (red in Fig. 5) were averaged together. Then, each data set was assigned to the group for which the predictions indicated the highest probability (winner-takes-all). This allowed us to determine the number of correct classifications and the number of specific confusions between classes (Fig. 6).

**Figure 6.**
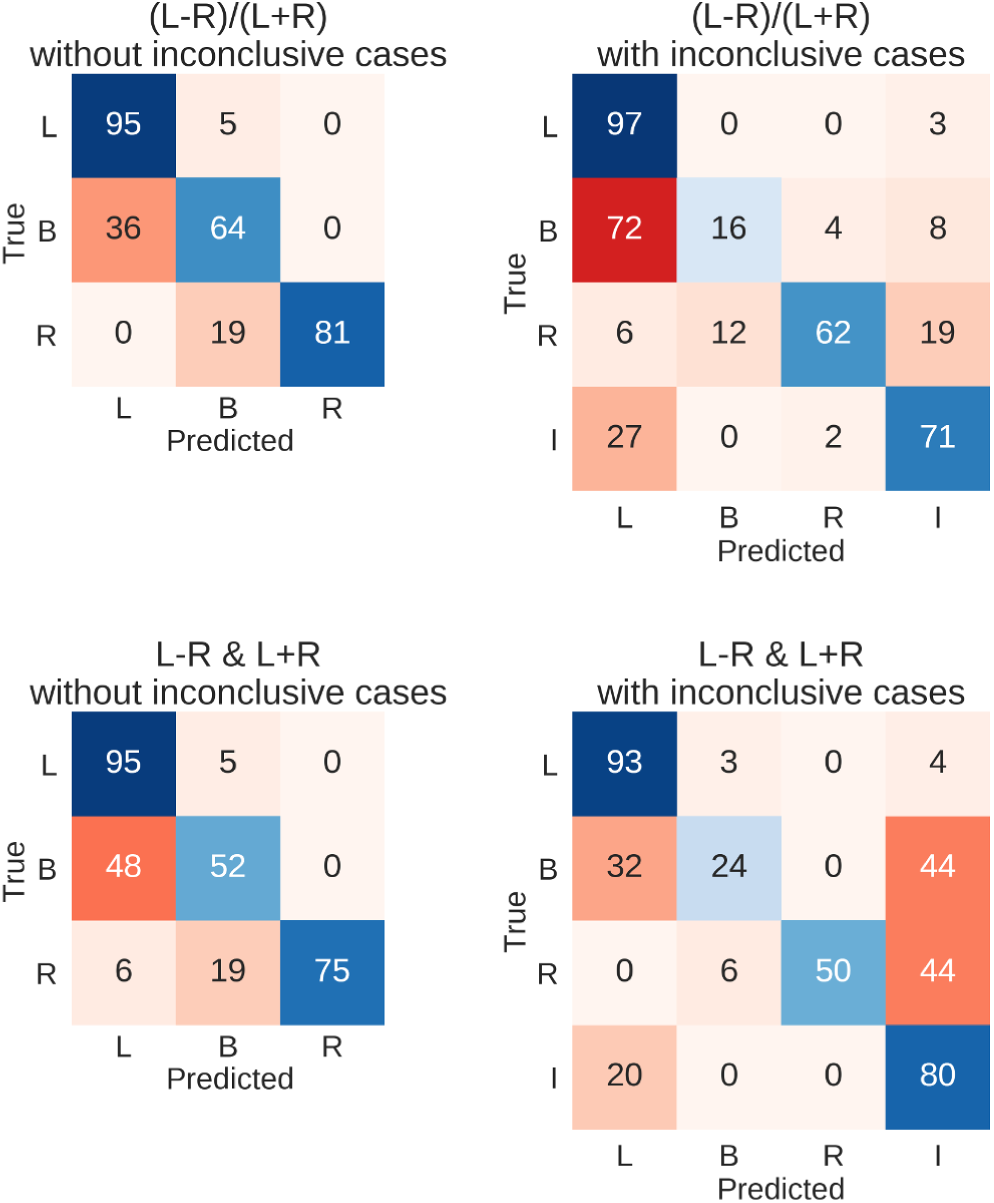
Confusion matrices for the four types of prediction. Results for predicting the test data set using held-out 262 fMRIs data sets are shown. Either the LI or its deconstruction into laterality and strength was used, and inconclusive cases were either excluded or included in the sample. Each row of the matrix sums up to 100% (save rounding errors), as it represents the true cases. The columns represent the predictions made. Correct predictions are plotted along the diagonal in blue. Confusions are plotted off-diagonal in red. Abbreviations: L=left; B=bilateral; R=right; I=inconclusive.

The resulting overall accuracies for the test data were almost identical to the top accuracies from the training data set. For the common LI, the held-out data were classified correctly in 91% of the cases when only conclusive data were used and in 82% of cases when inconclusive data were included. Accuracies for when the LI was deconstructed into laterality and strength reached 89% and 82%, respectively. Fig. 6 further illustrates the results with the diagonal showing correct predictions and the off-diagonal cells showing confusions. When only conclusive cases were included, both approaches showed similar patterns of hits and confusions with high accuracies for the left- and right-lateralised cases and frequent confusions of bilateral cases with left-lateralised cases. When inconclusive cases were added, the two approaches seemed to diverge more; the LI still confused bilateral cases with left-lateralised cases. On the other hand, the two-dimensional approach tended to mistake bilateral and right-lateralised cases for inconclusive cases more often.

### 3.4 Visualisation of all laterality scores

To better understand the way the different approaches reached their maximum accuracies, we plotted the distribution of data for the best parameter combination of each approach. Here, each patient’s laterality score is the average of all scores from the above-chance parameters. Fig. 7 shows that when only conclusive cases were included, there was a clear separation of classes using the LI. When inconclusive cases were included, the use of higher thresholds led to more extreme values (i.e., laterality scores approaching +1 and −1), especially for the left-lateralised cases, which showed overall more extreme scores. As the high thresholds (and the low denominators of the LI) led to more scores at the extremes and fewer around zero, the bilateral cases were pushed away from the middle of the scale. Finally, some cases were missing from the plot because at the high thresholds, the denominator of the LI was equal to zero, and no score could be computed.

**Figure 7.**
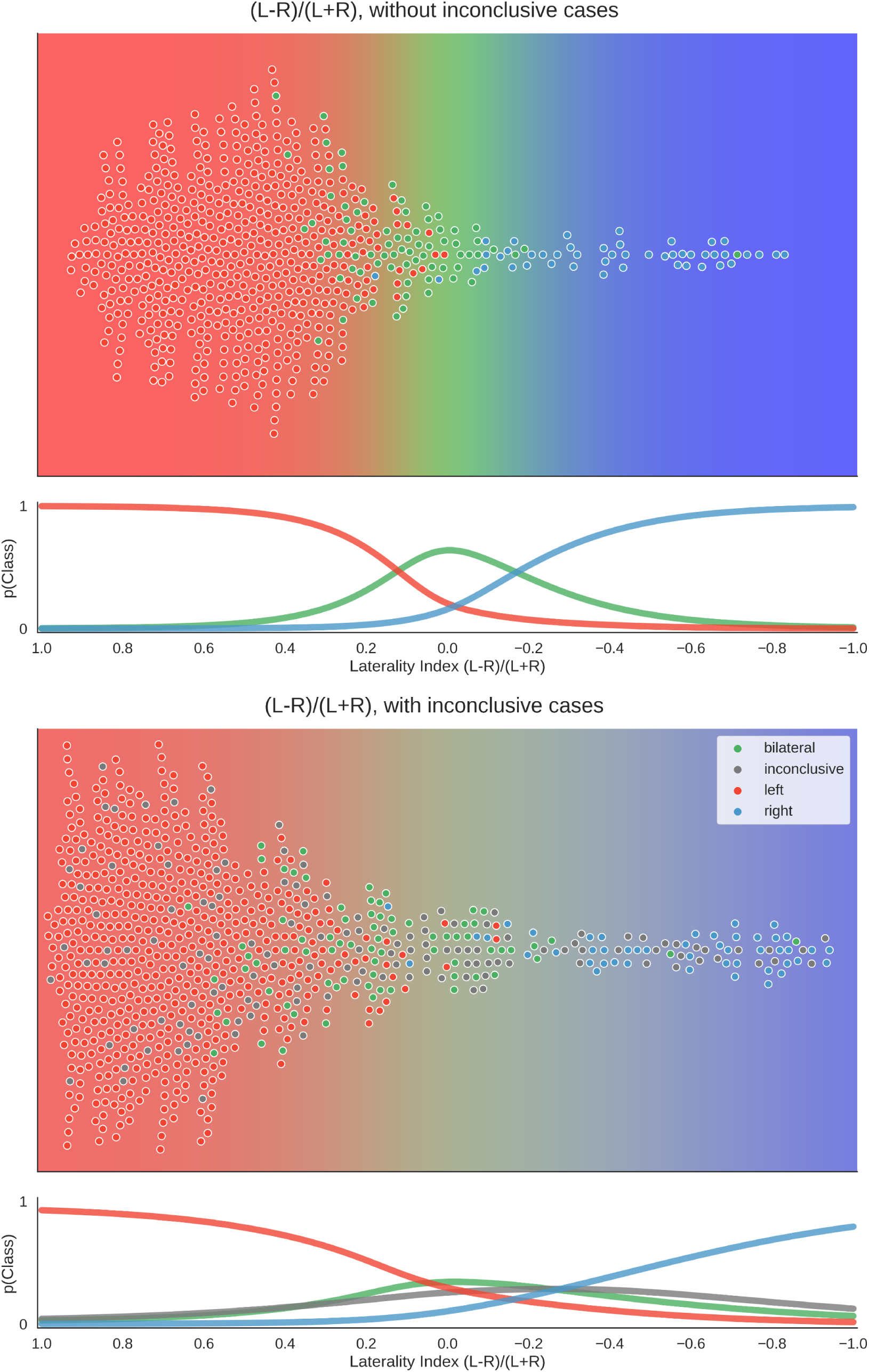
Distribution of data and predictions for top parameter combinations for the LI. The upper part of each plot shows the data as a distribution along the x-axis, with the background colour indicating the probabilistic prediction of the classifier. The human evaluation of each fMRI is indicated by the colour of the dots. The mean LI of each case on the axis was computed as the mean score from all above-chance parameter combinations (see Fig. 5). As positive LI scores are indicative of left-side dominance, the x-axis is inverted and goes from +1 to −1, so that the left side of the plot shows the left-lateralised cases. The lower panel of each plot shows the probabilities of belonging to each class along the range of LI values.

For the two-dimensional approach, the data were plotted with the numerator (L-R) on one axis and the denominator (L+R) on the other. Accordingly, in Fig. 8, each case has one mean score for each dimension. With conclusive data only, we see that the bilateral class spread out as strength increased. Thus, a case was more likely to be classified as bilateral when activation strength was high and more likely to be left- or right-lateralised when activation strength was low. This mimicked the one-dimensional LI, where a difference score based on low activation strength led to a strong laterality score (e.g., nine voxels in the right-hemisphere and one voxel in the left hemisphere is equal to −0.8), while the same absolute difference at a high level (e.g., 54 voxels in the right-hemisphere and 46 voxels in the left hemisphere is equal to −0.08) would be more indicative of bilaterality.

**Figure 8.**
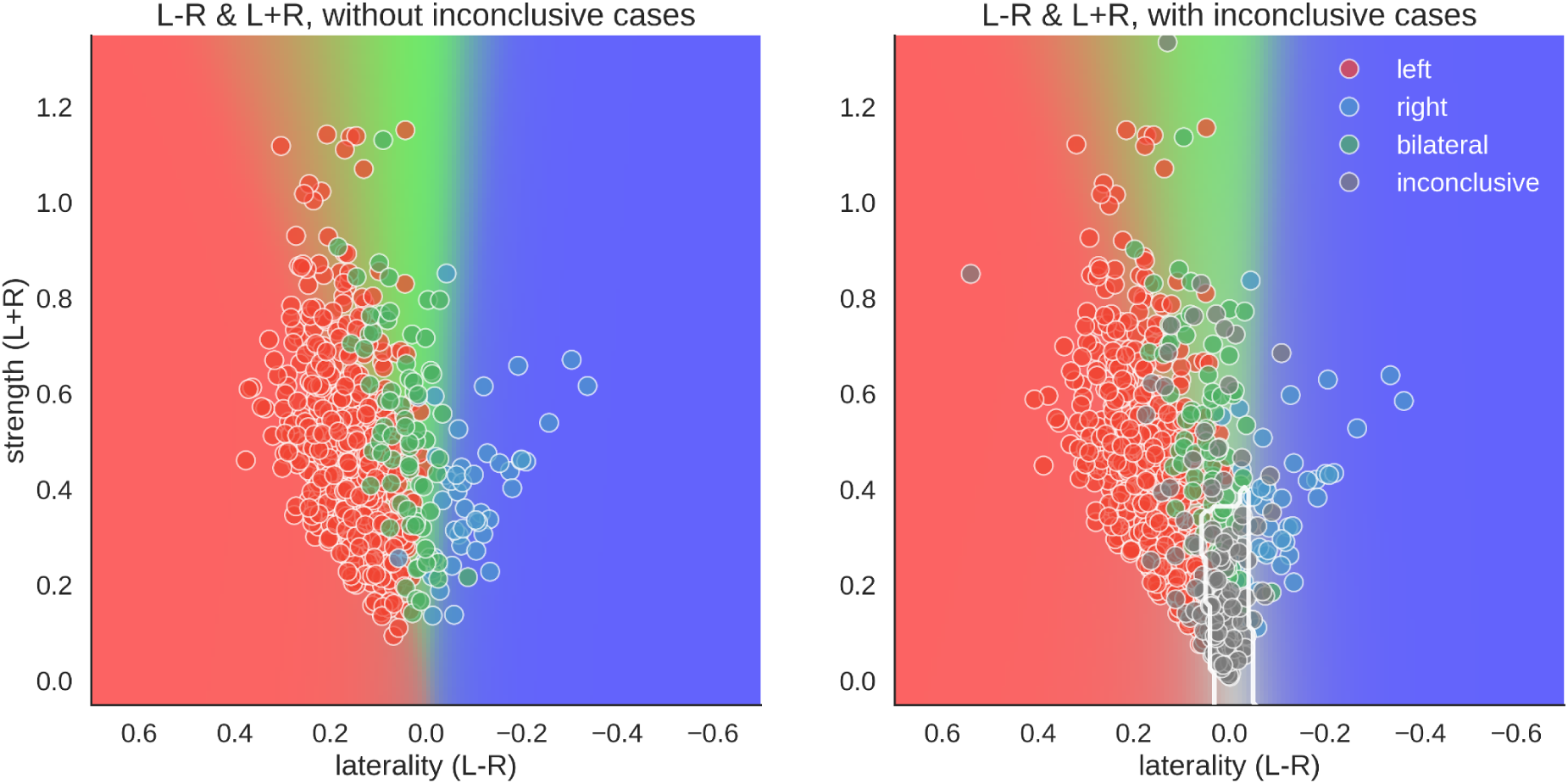
Distribution of data and predictions for top parameter combinations for the two-dimensional space. The x-axis of each plot shows laterality of the data (L-R), and the y-axis shows the overall activation (L+R). As positive LI scores are indicative of left-side dominance, the x-axis is inverted and goes from +1 to −1, so that the left side of the plot show the left-lateralised cases. The mean LI of each case on the axis was computed as the mean score from all above-chance parameter combinations in Figure 5. The background colour shows the average behaviour of the classifier combinations, as a colour-coded probability value. The boundary of the classifier which separates the inconclusive class from the rest is drawn as a white contour. The human evaluation of each fMRI is indicated by the colour of the dots.

However, when inconclusive cases were included, the two-dimensional approach differed more prominently from the LI. For the common LI, the inconclusive cases did not have predictable values, but in the two-dimensional approach, they occupy a specific part of the prediction space –namely, a case was classified as inconclusive if it had both low strength and low laterality values. In contrast, a case that also had low laterality values but showed high activity strength was classified as bilateral.

To illustrate how these analyses might be used on the level of individual patients, Fig. 9 illustrates the results of two cases.

**Figure 9.**
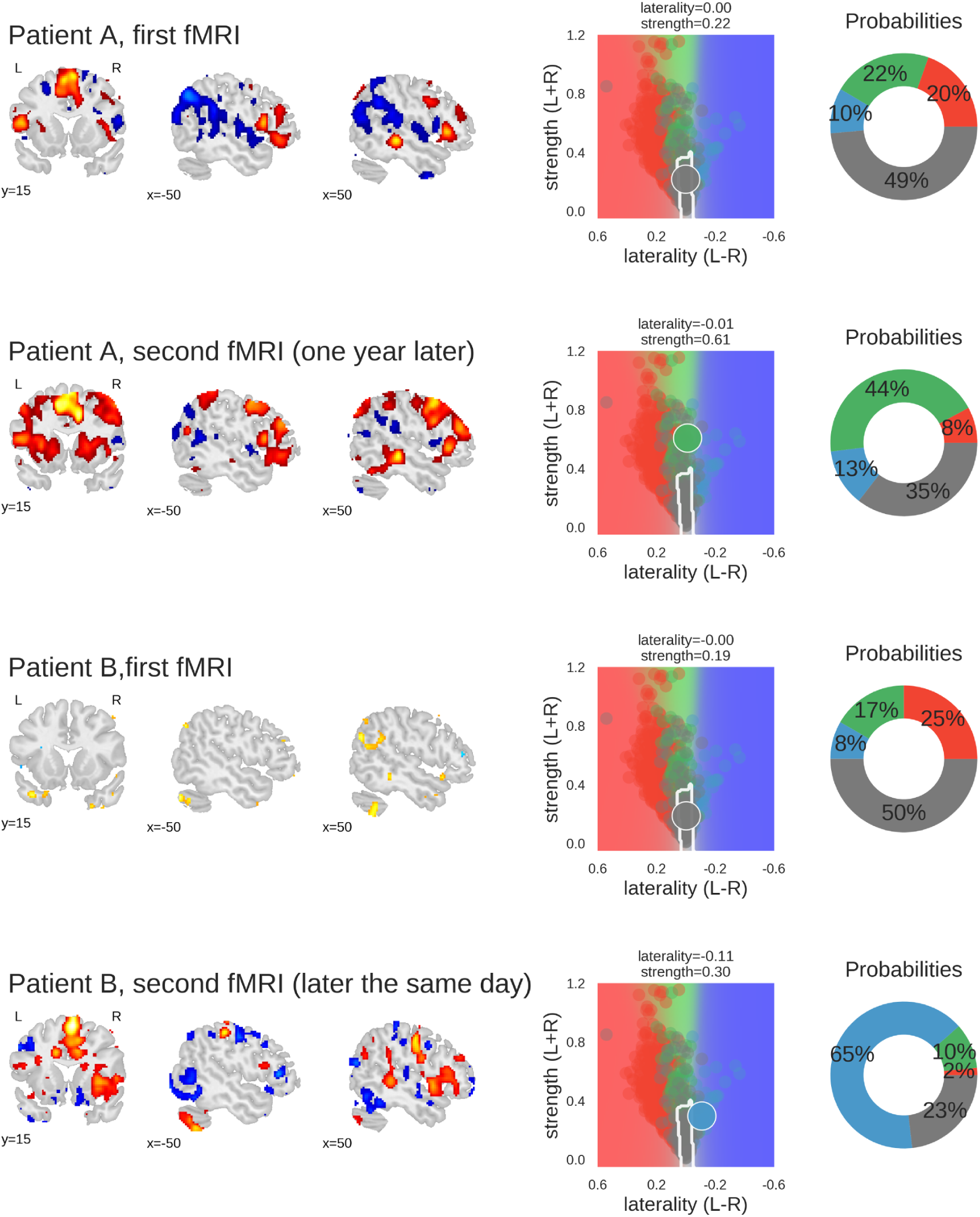
Examples of classifying individual fMRI data sets. For each fMRI, three sample slices of whole-brain activity are shown on the left hand (threshold at t=3); the middle plot shows the prediction space for the two-dimensional approach; the probabilities of belonging to each class are plotted as percentages on the right-hand plot. The four classes are represented by the colours red (left), green (bilateral), blue (right) and grey (inconclusive), as in the other figures. Patient A underwent fMRI language lateralisation twice, with the two measurements more than one year apart. Both times, the human evaluator assessed the activity pattern as bilateral, which was later confirmed by Wada-testing (bilateral Wada LI was −0.24). The 2d-method was too conservative the for the first measurement, but correctly classified the second fMRI (which showed stronger activity but no change in laterality) as bilateral. Patient B underwent fMRI language lateralisation twice on the same day, as the human evaluator assessed the first fMRI to be inconclusive. The second fMRI (which showed stronger activity and pronounced right-lateralisation) was assessed as being indicative of right-lateralised language. This is also captured by the predictions of the 2d-method. The patient later underwent Wada testing which confirmed the right-lateralised language (bilateral Wada LI was −1). Code used to generate the results is available at github.com/mwegrzyn/laterality-index-deconstruction.

### 3.5 Comparison with benchmark

To move beyond a comparison against guessing, we re-ran the above analyses using three variations of LI computations as implemented in the LI-Toolbox (Wilke and Lidzba, 2007) to train SVCs and then predict the language lateralisation determined by the human evaluator. To illustrate the behaviour of the bootstrap-LI (Wilke and Schmithorst, 2006), Fig. 10 shows the distribution of the LI values of all cases and the prediction space based on the training data. This follows the same rationale as described in section 3.4. The bootstrap-LI allowed to reach 92% correct predictions when only conclusive cases were included and 76% when inconclusive cases were also present. In the four-class case, there were more inconclusive cases with LIs around zero than there were bilateral cases, prohibiting the identification of bilaterality altogether.

**Figure 10.**
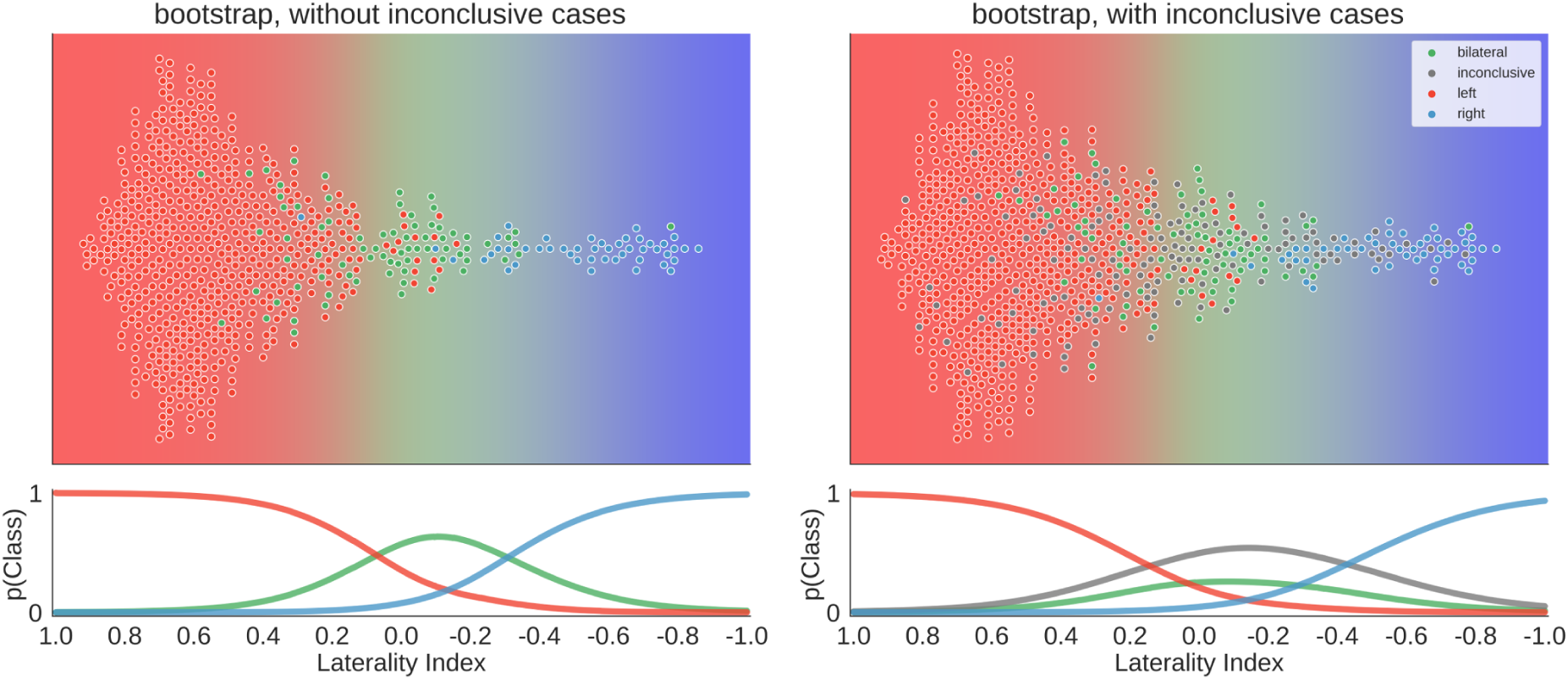
Distribution of data and predictions for the bootstrap-LI. The upper part of each plot shows the data as a distribution along the x-axis, with the background colour indicating the probabilistic prediction of the classifier. The human evaluation of each fMRI is indicated by the colour of the dots. The LI of each case on the axis was computed using the bootstrap-LI (Wilke and Schmithorst, 2006). As positive LI scores are indicative of left-side dominance, the x-axis is inverted and goes from +1 to −1, so that the left side of the plot shows the left-lateralised cases. The lower panel of each plot shows the probabilities of belonging to each class along the range of LI values.

Overall, the accuracies of all three approaches of the LI-Toolbox were comparable to our analyses (Fig. 11). Within the LI-Toolbox, there was no significant difference between adaptive, bootstrap, and fixed-threshold methods.

**Figure 11.**
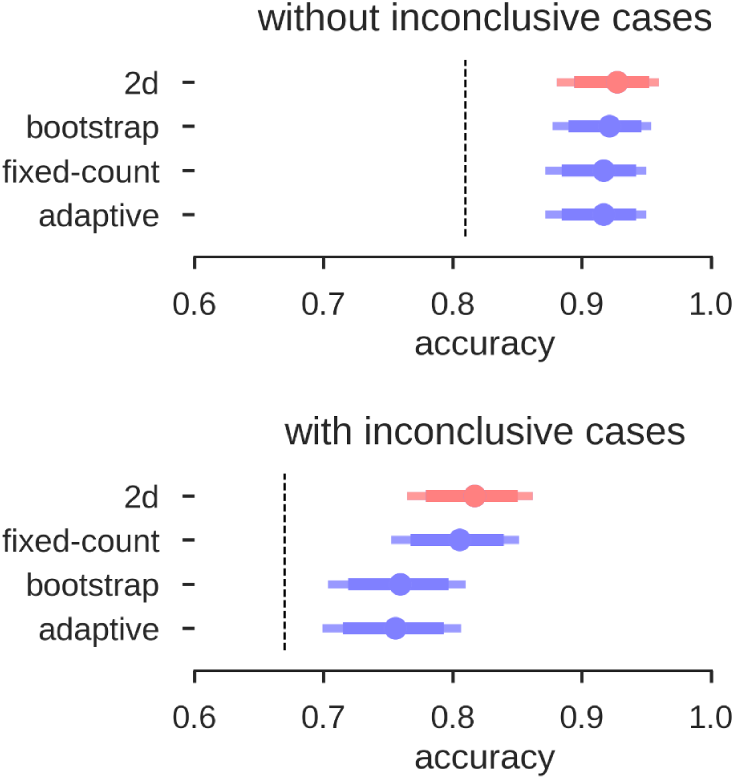
Accuracies for prediction of the test data. Mean accuracy (dot), the 84% confidence interval (thick line) and the 95% confidence interval (light line) are depicted; chance performance (always guessing left) is indicated by the dashed line. 2D: our approach with separate measures of laterality (L-R) and strength (L+R), highlighted in red; fixed-count: counting voxel at fixed threshold of t=3; adaptive: LI computation with thresholding at the mean intensity of the image; bootstrap: main output of the bootstrap-LI (Wilke and Schmithorst, 2006).

### 3.6 Validation with Wada test results

To validate all of the above analyses, we selected 80 fMRI data sets from 62 patients for which Wada test results were available. Instead of predicting the human evaluations (left, bilateral, right, inconclusive), we now tried to predict the Wada test result (left, bilateral, right). The same classifiers as above were used (i.e., derived from the training set with human evaluations as labels). No new or additional training on the Wada scores was performed.

The results for the different approaches are depicted in Fig. 12. When using the LI-Toolbox immediately, without preselecting the fMRI data regarding their conclusiveness, accuracies between 71% and 75% were reached. When using our two-dimensional approach to first exclude cases deemed inconclusive (34 fMRI data sets according to our approach) and then try to predict the Wada result of the remaining cases, a higher accuracy of 83% for the remaining 46 cases was reached. When combining the different approaches (i.e., ours to exclude inconclusive cases and the LI-Toolbox to classify the remaining ones), accuracies of 85% correct were reached. Accuracies for our 2D approach and 2D-LI hybrids were all significantly above 63% chance. That the pre-selection of inconclusive data contributed to increasing the accuracy is illustrated at the bottom of Fig. 12. There, only the Wada test results of cases deemed inconclusive by the human evaluator were predicted. Our approach excluded 20 of the 21 inconclusive cases and made a correct prediction for the one remaining case, while the adaptive LI was significantly below guessing (always guessing left, indicated by the dashed line, being the superior strategy).

**Figure 12.**
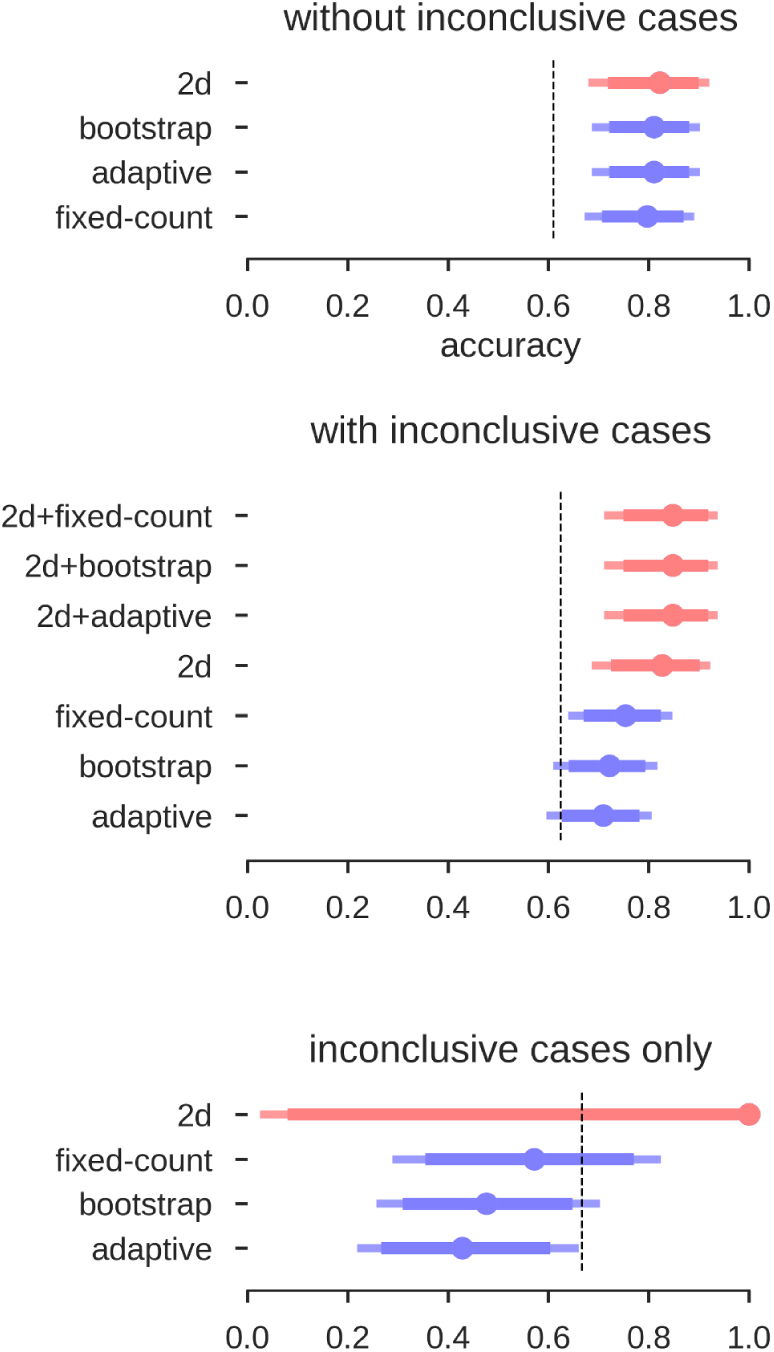
Accuracies for the prediction of Wada test results. Mean accuracy (dot), the 84% confidence interval (thick line) and the 95% confidence interval (light line) are depicted; chance performance (always guessing left) is indicated by the dashed line. 2D: our approach with separate measures of laterality (L-R) and strength (L+R), highlighted in red; fixed-count: counting voxel at fixed threshold of t=3; adaptive: LI computation with thresholding at the mean intensity of the image; bootstrap: main output of the bootstrap-LI (Wilke and Schmithorst, 2006). ‘+’ sign indicates whenever the 2D approach was used to exclude inconclusive cases and another LI-measure when then used to classify the remaining cases (highlighted in red).

## Discussion

In the present study, we used data from a large sample of epilepsy patients performing an fMRI verbal fluency task. We aimed to evaluate how well an experienced human evaluator’s assessment of language lateralisation, based on free visual inspection of whole-brain fMRI patterns, can be predicted from an LI value.

When using high-quality data (i.e., excluding inconclusive cases based on low fMRI activity), above-chance accuracies of up to 92% correct classifications were reached. This is in accordance with previous studies on language lateralisation based on conclusive fMRIs, indicating that estimating laterality from high-quality fMRI activity data is robust (Woermann et al., 2003; Jones et al., 2011; Janecek et al., 2013). Given the large amount of data reduction that goes into computing an LI, this level of accuracy is noteworthy. When including inconclusive data, and thereby more closely simulating the clinical context (in which noisy data are not unusual), the common LI also produced high accuracies and was as good as a two-dimensional approach that allows grouping the data both by laterality (L-R) and by strength (L+R). That increasing the dimensionality of the data does not allow for better classification indicates that the common LI is a useful method of data reduction despite its flaws (see Jansen et al. 2006; Seghier 2008). While both approaches perform equally well at the group level, the two-dimensional approach has the advantage that it can always be evaluated, as we never have to divide by zero. This might be a more desirable approach in the clinical context compared to using the heuristic that whenever the LI yields an error, we assume that laterality cannot be determined. While inconclusive cases scatter along the whole continuum of the common LI (from −1 to +1), they occupy a predictable range of values in the two-dimensional space. There, inconclusive cases scatter around zero on the L-R scale, given that they contain little information about laterality; and because they show little language-related activity on either side, they scatter around zero on the L+R scale (see the grey dots in Fig. 1 and Fig. 8). Of note, our two-dimensional approach is merely a deconstruction of the common LI formula into its numerator and denominator. Therefore, it does not require abandoning the LI for a new approach or combining multiple methods but merely taking advantage of the information already contained in the LI. That it provides a predictable range into which inconclusive cases will fall also sets this approach apart from the established adaptive and bootstrap methods for LI computation (Wilke and Schmithorst, 2006; Wilke and Lidzba, 2007). While fixed threshold methods have the problem that low activity data produce LIs close to the extremes (+1 and −1), our results show that for adaptive methods, low activity data produce LIs close to zero. This can make it especially challenging to correctly identify bilateral cases. However, these results must be interpreted with caution, as it is also possible that all of the inconclusive cases with bootstrap-LI values around zero were truly bilateral patients. In this case, this LI method would be unfairly penalised for out-performing the human evaluations, which assigned these data sets to the inconclusive class. By using the Wada test results as the gold standard of language lateralisation, we were able to show that no method performed above chance for the group of inconclusive cases. This strengthens the notion that modelling an inconclusive class for which no reliable prediction can be made is a useful strategy to reduce misclassifications. While it is costly to discard data, recognising that a case cannot be evaluated with sufficient certainty might still pay off, for example by allowing a more targeted repetition of measurements. This would also take advantage of the non-invasiveness of fMRI, which sets it apart from the Wada test.

The validation of our approach using information from the Wada test is also important because we trained our predictions using only human evaluations of the fMRI data. Using human evaluations is problematic because they are based on the fMRI data themselves and not a truly independent criterion. Also, they cannot replace the Wada test as the gold standard for validation. However, we know that the agreement of human evaluation of fMRI and Wada test results is very high (Woermann et al., 2003). Therefore, using human evaluations as the criterion allowed us to assemble very large samples (compared to the Wada test, which is much less frequently performed), making data-driven methods more feasible. Hence, the approach of the present study would not have been possible if only one or a handful of atypical cases were available, as is frequently the case in the literature (see the review by Dym et al. 2011). Despite never using Wada test data during training, we were still able to predict the Wada results with accuracies significantly above chance. This lends support to the chosen approach and confirms the high concordance of fMRI and Wada test results (Binder, 2011; Szaflarski et al., 2017). Given that the sample of Wada test patients contained a large proportion of atypical patients (37%) and that Wada tests are usually administered in difficult cases, accuracies around 80% are noteworthy.

When comparing the accuracies between our approach and the different variations implemented in the LI-Toolbox (Wilke and Lidzba, 2007), we saw that the LI is very robust across its different implementations. Although we had a large sample of validation data, we were not able to find systematic differences between fixed-threshold and adaptive or bootstrap methods. Meanwhile, it is possible that combining a fixed-threshold LI with an adaptive LI might be superior to using one or the other by itself. To recognise inconclusive cases, a fixed threshold seems ideal. Only when the threshold is kept fixed across patients, differences in the denominator of the LI formula (L+R) can be used to compare activity strength. If an adaptive method keeps the denominator’s value constant (e.g., L+R must always equal 50% of the ROI size; Wilke and Lidzba 2007), this differential information will be lost. Afterwards, if a data set is deemed fit for further analysis, an adaptive threshold might be optimal because it maximises the variance in the numerator of the LI formula (L-R) by keeping the denominator constant.

While assigning a measure of uncertainty to each individual prediction is useful to reduce mistakes, future studies should explore whether data from an inconclusive fMRI can somehow be salvaged to correctly predict the patient’s Wada test result. While the current study’s incorporation of inconclusive class allowed testing predictions in a realistic context, classifying a case as inconclusive cannot be the ultimate goal of diagnostics. Alternative data analysis techniques such as pattern analysis methods (Zago et al., 2017) might be used to take full advantage of the data and successfully predict the true type of lateralisation of each patient.

Furthermore, none of the approaches presented here showed a satisfying sensitivity regarding the detection of bilateral cases (see Fig. 6). This might reflect that many instances of bilaterality cannot be well expressed with a simple LI. For example, crossed lateralisations with left-sided activity in Broca’s area and right-sided activity in Wernicke’s area (Kurthen et al., 1992) might by definition be unsuitable to be represented by a simple score based on one ROI. Multiple ROIs might have to be considered simultaneously (Benjamin et al., 2017) to improve the characterisation of each patient’s unique type of language lateralisation. Also, some studies have indicated that additional classes of language lateralisation might be needed to better understand the different subtypes of atypicality (Berl et al., 2014).

While the usefulness of fMRI for language lateralisation in epilepsy has substantially improved since the inception of the method (Szaflarski et al., 2017), it still needs more validation work to make it feasible for reliably predicting language lateralisation (i) in single cases and (ii) in a clinical setting. Although much research has focused on how to compute the LI, it is less well understood how an LI value should be best translated into a categorical decision. Apart from the LI threshold, brain regions for data extraction and cutoffs for grouping the LI values into clinical categories must be chosen. Also, it must be considered that some data sets simply do not contain enough diagnostic information to allow for a confident decision. In our study, we tried to justify every step of such a decision-making process, using data-driven methods throughout. The validation of our approach shows that the LI is very robust across its different implementations. It can predict human evaluations of language lateralisation as well as Wada test results with substantial above-chance accuracies. Our results indicate that by taking advantage of all information contained within the LI, its clinical utility could be further improved.

## Acknowledgments

Kirsten Labudda holds a Junior-Professorship at Bielefeld University endowed by the von Bodelschwinghsche Stiftungen Bethel. The funding source had no influence on the study’s design, data collection, analyses, interpretation, manuscript preparation and submission.

We would like to thank Anna Doll for help with data collection.

## Data Availability

fMRI group result maps are available on NeuroVault

neuro neurovault.org/collections/3887/

Code to reproduce the results and figures, as well as to recreate this manuscript, can be found on GitHub

code github.com/mwegrzyn/laterality-index-deconstruction

